# Dissection of the Global Responses of Mandarin Fish Pyloric Caecum to An Acute Ranavirus (MRV) Infection Reveals the Formation of Serositis and Then Ascites

**DOI:** 10.1101/2025.01.02.631049

**Authors:** Wenfeng Zhang, Yong Li, Xiaosi Wu, Qianqian Sun, Yuting Fu, Shaoping Weng, Jianguo He, Chuanfu Dong

## Abstract

Mandarin fish ranavirus (MRV), a variant of largemouth bass virus (LMBV), is a distinct member of the genus *Ranavirus* within the family *Iridoviridae*. Acute MRV infection predominantly affects the pyloric caecum, a critical visceral organ in mandarin fish, and is hypothesized to drive the characteristic external clinical sign of severe ascites. In this study, we reveal that acute MRV infection initially targets the serosal layer of the pyloric caecum of mandarin fish, leading to rapid progression into fibrinous serositis characterized by serosal hypertrophy, fibrosis, hyperemia, edema, and tissue adhesions. Using single-cell RNA sequencing, we dissect the cellular composition of epithelial, immune, and stromal populations, identifying significant enrichment of macrophages and granulocytes, alongside T and natural killer cells, as key mediators of acute cytokine and inflammatory responses. Then, robust experimental evidence demonstrates that MRV attacks specific immune cell subsets of T and B cells and stromal cells of fibroblasts, myofibroblasts, endothelial cells, and pericytes, resulting in upregulation of genes and pathways associated with extracellular matrix (ECM) formation, collagen biosynthesis, and vascular remodeling in the hyperplastic serosal zone. Additionally, both host-derived type V collagens and MRV-encoded collagens are implicated in ECM formation in the hypertrophic serosa. Collectively, this study provides a comprehensive single-cell resolution analysis of the pyloric caecum’s response to acute MRV infection and highlights virus-driven serositis as the underlying cause of severe ascites in mandarin fish.

**Importance:** The pyloric caecum is a vital digestive and immune organ in many bony fish species, including the mandarin fish, a carnivorous species with an exceptionally developed pyloric caecum comprising 207–326 caeca per individual. While MRV/LMBV infects various fish species, severe ascites is uniquely observed in infected mandarin fish. This study demonstrates that acute MRV infection induces fibrinous serositis in the pyloric caecum, characterized by hyperemia, edema, and hyperplasia, ultimately resulting in ascites and mortality. Leveraging single-cell RNA sequencing, we provide a detailed landscape of the cell types affected or involved in the inflammatory response, revealing their roles in the pathogenesis of serositis. These findings advance our understanding of MRV-induced pathology and its species-specific manifestations.

## Introduction

Iridoviruses are large, icosahedral viruses possessing a single molecule of double-stranded DNA (dsDNA). Their genomes range from 99 kilobases (kb) to 220 kb, encoding 97 to 221 open reading frames (ORFs) [1, 2]. According to the latest updated version of the International Committee on Taxonomy of Viruses (https://talk.ictvonline.org/taxonomy/), the family *Iridoviridae* comprises of seven genera. Among these, four genera (*Chloriridovirus, Iridovirus, Decapodiridovirus, and Daphniairidovirus*) infect invertebrates such as insects, crustaceans, as well as mollusks, while the remaining three genera (*Ranavirus, Lymphocystivirus*, and *Megalocytivirus*) infect a variety of cold-blooded vertebrates [3]. Among vertebrate-infecting iridoviruses, ranavirus stands out for its broad host range, infecting bony fish, amphibians, and reptiles. Ranavirus infections often cause high morbidity and mortality in commercially valuable fish, amphibians, and threatened wildlife [4–8].

The clinical symptoms of ranavirus infections are generally systemic, characterized externally by multifocal cutaneous hemorrhages and erythema, and internally by necrosis of hematopoietic tissues, vascular endothelium, and epithelial cells, along with hemorrhage and the formation of intracytoplasmic basophilic inclusion bodies [9–12]. The affected organs often include the liver, spleen, kidney (pronephros and mesonephros), and occasionally the gastrointestinal mucosa, lymphoid tissue, and neuroepithelial tissue [4, 6, 11–14]. Mandarin fish ranavirus (MRV), also known as a variant of largemouth bass virus (LMBV), represents a highly distinctive member in genus *Ranavirus* [2]. The clinical symptoms of LMBV infection are generally like those of other ranaviral infections. Under laboratory conditions, the featured external sign of LMBV-infected either largemouth bass (*Micropterus salmoides)* or smallmouth bass *(M. dolomieu)* was the development of ulcerative dermatitis and necrotizing myositis in the site of infection [13, 15, 16]. By contrast, the acute MRV infections in mandarin fish (*Siniperca chuatsi*) are marked by severe ascites, a unique external clinical feature not observed in other *Ranavirus* infections [17, 18].

A recent histopathological investigation showed that pyloric caecum of mandarin fish was the most attacked visceral organs upon acute MRV infection and then proposed that MRV might be a digestive tract pathogen to mandarin fish, which might contribute to the featured ascites syndrome [19]. The pyloric caecum locates in the second segment of the gastrointestinal tract, serves critical roles in digestion, nutrient absorption, and immune responses in bony fish [20]. While Aristotle firstly described the pyloric caecum as early as in 345 BCE [21, 22], its precise functions have been historically debated. Modern studies have clarified its roles in food storage, digestion, and as an active immune organ [22–24]. Mandarin fish possess a highly developed pyloric caecum, with each individual hosting 207–326 caeca [25], a stark contrast to the 25 caeca observed in largemouth bass [23]. Our recent study showed that the pyloric caecum of mandarin fish is the primary targeted internal organ upon acute MRV infection, which has not been documented in other ranaviral infections [19]. By contrast, in LMBV-infected largemouth bass, the over-inflated swim bladder with thick, yellow or brown exudate was descripted as one of the most featured internal clinical signs [26]. Despite these observations, the mechanisms underlying MRV-induced ascites and the contrasting organotropism of MRV and LMBV remain poorly understood.

Single-cell RNA sequencing (scRNA-seq) has transformed our understanding of cellular heterogeneity in various fields, including immunology, oncology, and virology [27–30]. This technique enables high-resolution analysis of transcriptional states at the single-cell level, offering unique insights into viral tropism, host cell responses, and virus-host interactions [31–37]. In this study, we revisited the histopathological changes in the pyloric caecum of mandarin fish following acute MRV infection and applied scRNA-seq to elucidate the transcriptional landscapes of infected and uninfected pyloric caecum cells. Our findings identified targeted immune (B and T lymphocytes) and stromal cells (fibroblasts, myofibroblasts, endothelial cells, and pericytes) as key contributors to the inflammatory cytokine response and the development of severe fibrinous serositis. Together, these processes drive ascites syndrome and mortality in infected mandarin fish.

## Results

### Pyloric caeca serve as the most attacked internal tissue, and the serosa layer bears the primary infection

Through a series of experiments including necropsy observation, real-time quantitative polymerase chain reaction (RT-qPCR), as well as immunohistochemistry (IHC) and immunofluorescence (IF), we systematically examined the infection sites and replication patterns of the acute MRV infection (**Fig. 1** and **Fig. S1**). The results revealed that the MRV infection caused severe congestion and edema in the pyloric caeca, hightlighting its high susceptibility to the virus (**Fig. 1A**). Both RT-qPCR and IHC analysis showed significantly higher viral loads and stronger MRV signals in the pyloric caeca compared to other tissues (**Fig. 1B and C**). Microscopically, the health pyloric caecum displays a layered structure comprising of mucosal, submucosal, mucosal muscle, and the outer serosal layer (**Fig. 1D and F**). In contrast, the transverse sections of infected pyloric caeca, stained with anti-MRV monoclonal antibodies, showed a selective highly concentrated of MRV-positive signal in the serosal structures by IF (**Fig. 1E**). Few MRV positive signals were detected in the mucosal, submucosal, or mucosal muscle layers, indicating a specific tropism of MRV to the serosa. Furthermore, in contrast to the uninfected healthy pyloric caeca, the infected pyloric caeca revealed significant expansion of the serosal structure, providing a favorable environment for extensive viral replication (**Fig. 1F**). A temporal analysis of the infection revealed a progressive formation of proliferative serosa with MRV infection (**Fig. S1**). At 1day post infection (dpi), slight hyperplasia was observed along the serosal layer. By 3 dpi, obvious hyperplasia filled the major gaps among pyloric caeca, and by 5 dpi, severe hyperplasia had completely filled these gaps. IHC and IF analyses confirmed that the serosal layer bears the primary MRV infection as early as 1 dpi, with high concentrations of MRV detected at 3 and 5 dpi, especially in the hypertrophic regions of infected pyloric caeca. In contrast, the interface of uninfected, healthy pyloric caeca remained clear and intact, with no hyperplasia or adhesion. All these results revealed that MRV infection in mandarin fish shows a unique tropism towards the pyloric caeca, among which, the serosal structure bears the primary infection of MRV.

**Fig 1.**
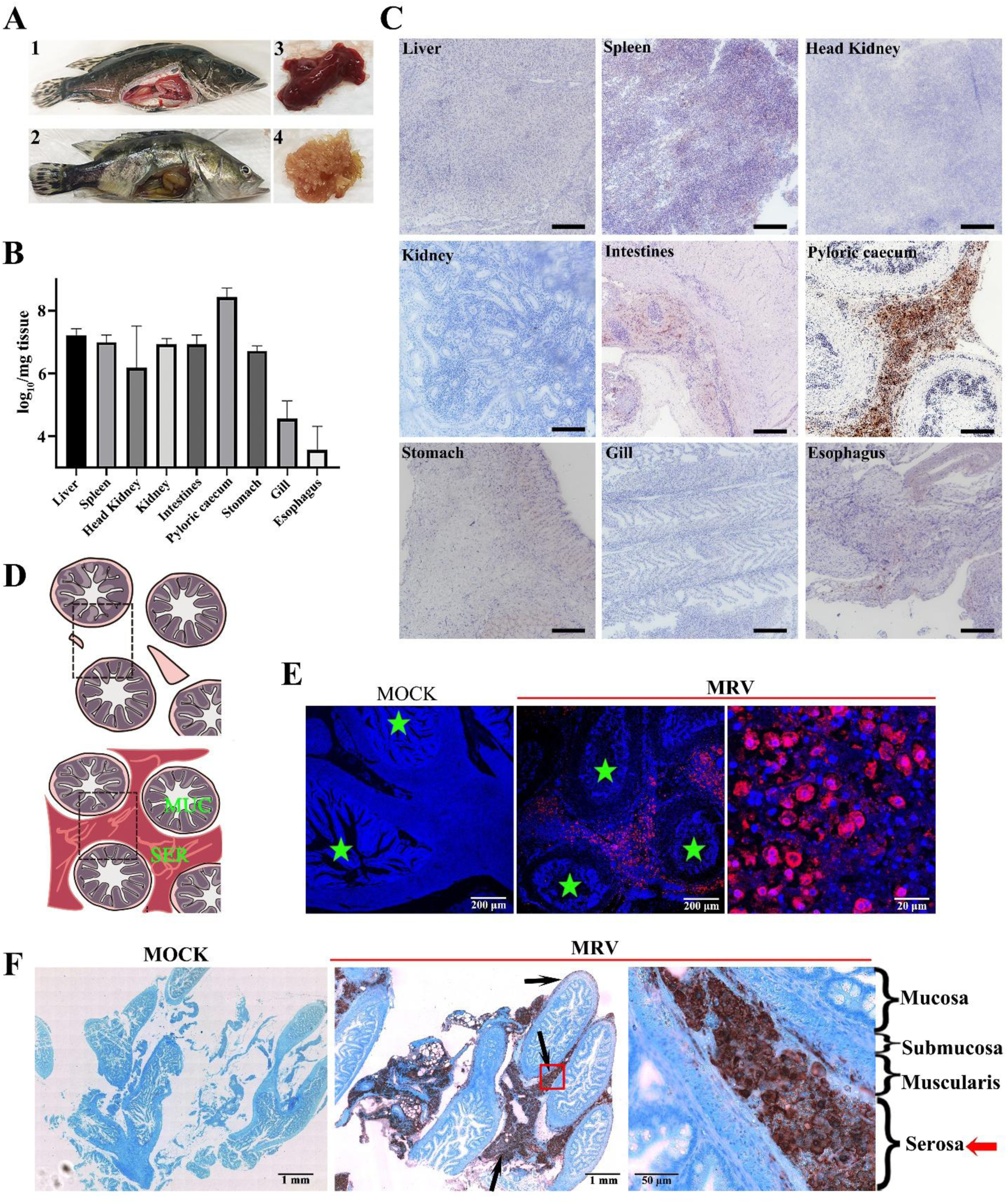
The main attack site of MRV in mandarin fish pyloric caeca. (A) The autopsy of MRV-infected mandarin fish. (A1) and (A3), the diseased fish was characterized by severe haemorrhage in pyloric caeca. (A2) healthy fish with normal pyloric caeca (A4). (B) Tissue distributions of MRV in various tissues by absolute RT-qPCR (n = 3). The highest viral load was determined in infected pyloric caeca. (C) Localization of MRV by immunohistochemistry (IHC). The strongest MRV-positive signals were found in infected pyloric caeca. Scale bars, 200 μm. (D) Schematic drawing of cross section of mandarin fish pyloric caeca. MUC, mucosa; SER, serosa. (E) Histo-immunofluorescence (HIF) of mandarin fish pyloric caeca infected with MRV. Numerous red fluorescence signals indicate MRV-infected cells by anti-MRV mAb 1C4. The asterisk indicates the digestive cavity. Scale bars of healthy fish and infected fish (left), 200 μm. Scale bars of infected fish (right), 20 μm. (F) Immuno-histochemistry (IHC) staining of MRV-infected pyloric caeca. Arrows indicate the sites stained by anti-MRV mAb 1C4. Serosa layer was observed as the most attacked site in pyloric caecum. Scale bars of healthy and infected fish (left), 1 mm. Scale bars of infected fish (right), 50 μm.

### Overview of the cell types of pyloric caeca by scRNA-seq

To comprehensively understand the cellular heterogeneity of pyloric caeca, scRNA-seq was performed on two uninfected (control) and two MRV-infected mandarin fish (**Fig. 2A**). To ensure robust data analysis, cells with fewer than 400 detected genes or more than 25% mitochondrial unique molecular identifier (UMI) counts were excluded. After eliminating data outliers based on the number of reads per cell (nUMI), the number of genes detected per cell (nGene), and thresholds for mitochondrial RNA genes, a total of 4,807 and 10,777 pyloric caecum cells were obtained from the control (C) and MRV-infected (MRV) mandarin fish, respectively (**Fig. S2**). To explore the cellular landscape, we applied T-distributed random neighbor Embedding (tSNE) reduction and unsupervised cell clustering, leading to the identification of eighteen cell clusters (**Fig. 2B**). Classification based on unique transcription profiles and the top five expressed genes in each group allowed us to categorize these clusters into epithelial cells, immune cells, stromal cells, and red blood cells, respectively. Utilizing well-established marker genes (**Fig 2E and F and Fig. S3**), we classified two major epithelial cell types (EPCAM) as ciliated epithelial cells (expressing CXCR4B, CCL25a) and secreting cells (expressing CHIA1, PYYA and TRYPSIN). Within the immune cell population, five distinct cell types were specified: T/NK cells (expressing TCRalpha, CD247, CD3delta), granulocytes (expressing MMP9 and NCF1), macrophages (expressing CSF1RB, MRC1 and MARCO), B cells (expressing IgM, Pax5, and CD79), and monocytes (expressing GRNA, MCP-1). Furthermore, three stromal cell types were also identified: fibroblasts (expressing FGFR2, FN1B), endothelial cells (expressing CDH5, CLDN7), myofibroblast and smooth muscle cells (expressing ACTA2 and MYH11). Notably, red blood cells (expressing HBA, HBB) were abundantly present in pyloric caecum of MRV-infected mandarin fish (**Fig. 2B**). With infection of MRV, the number and proportion of each cell subpopulation changed as shown in Fig. 2C and D.

**Fig 2.**
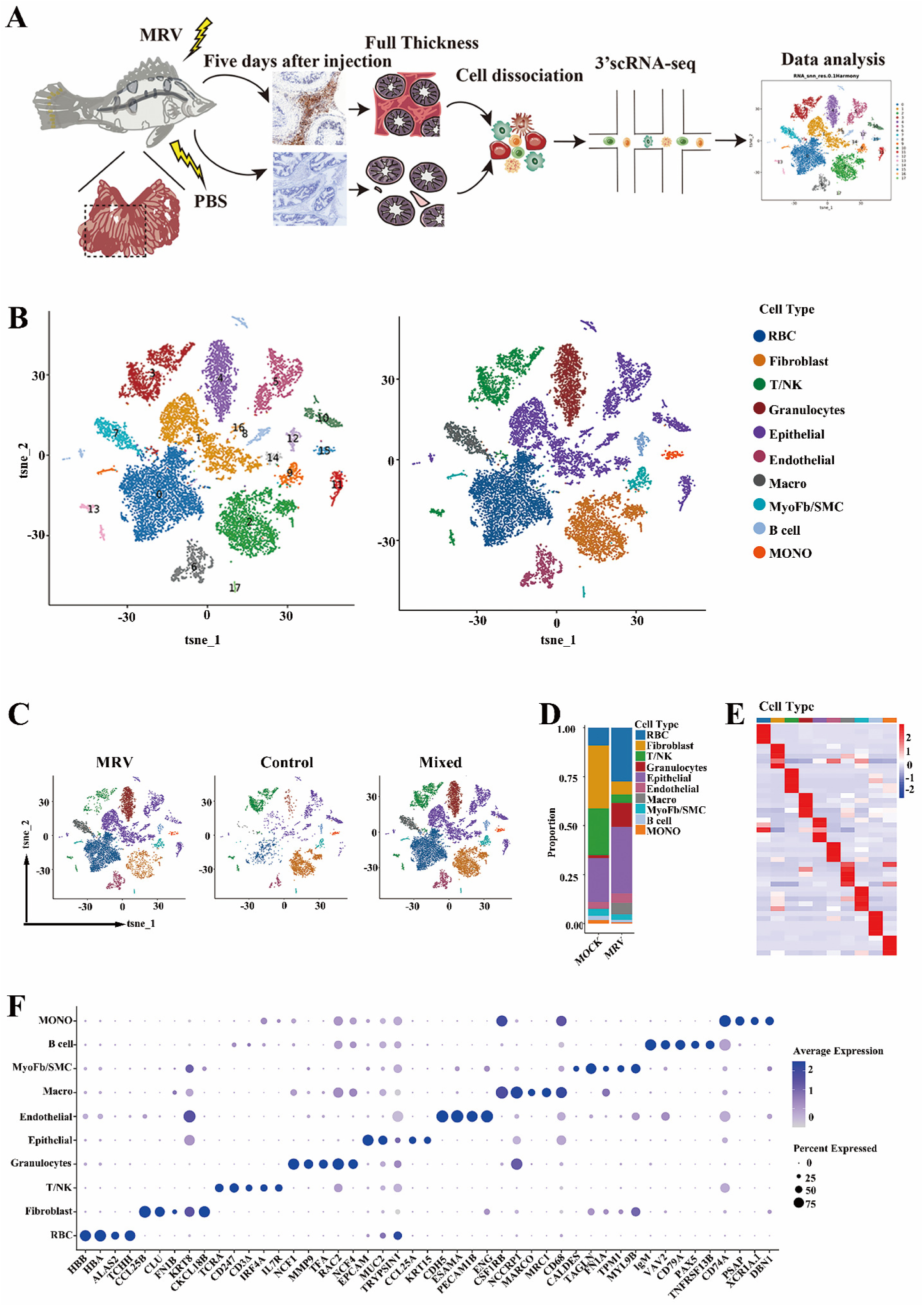
Cell sorting and categorization of cell types of mandarin fish pyloric caeca. (A) Overall strategy for cell sorting and single-cell data analysis. (B) The tSNE plot clustering of cells (left): different cell clusters are color coded; The tSNE plot showed the expression of pan marker genes in distinct cell clusters (right), and the gene expression level is color coded. (C) The tSNE plots align the clusters between control and MRV-infected mandarin fish. (D) Bar plots showing the proportion of each cluster in control and MRV-infected groups. (E) and (F) Average expression level and prevalence of selected major markers used to annotate the major cell types. Heatmap (E) and dotplot (F) of the expression of the marker genes in each cell type.

### Identification and characterization of epithelial subtypes and stromal subtypes

By re-clustering approximately 4,500 epithelial cells identified in the global analysis, twelve distinct molecularly defined subtypes were identified (**Fig. 3A**). The gene expression of tSNE plots maps were generated to identify marker genes for each epithelial subpopulation. Among these, non-secretory epithelial cells were designated Epithelial-1 to Epithelial-5, secretory cells are designated as Secretory-1 to Secretory-3, and goblet cells based on isotype-specific RNA markers. Additionally, two enteric nervous system cell subtypes were designated Neurocyte-1 and Neurocyte-2 (**Fig. 3B and S4A**). Given that minimal MRV positive signals were observed in mucosa of pyloric caecum cavity (**Fig. 1 and 2**), epithelial cells in the pyloric caeca were excluded as MRV target cells. Although MRV did not infect the epithelial layer of the pyloric caecum and the villi remained structurally intact, partial adhesion, epithelial exfoliation, and infiltration of inflammatory cells into the mucosal muscle were observed. These findings suggest that MRV infection may induce mucosal damage in the pyloric caeca, affecting both epithelial cell injury and subsequent recovery via cell proliferation (**Fig. 3C**).

**Fig 3.**
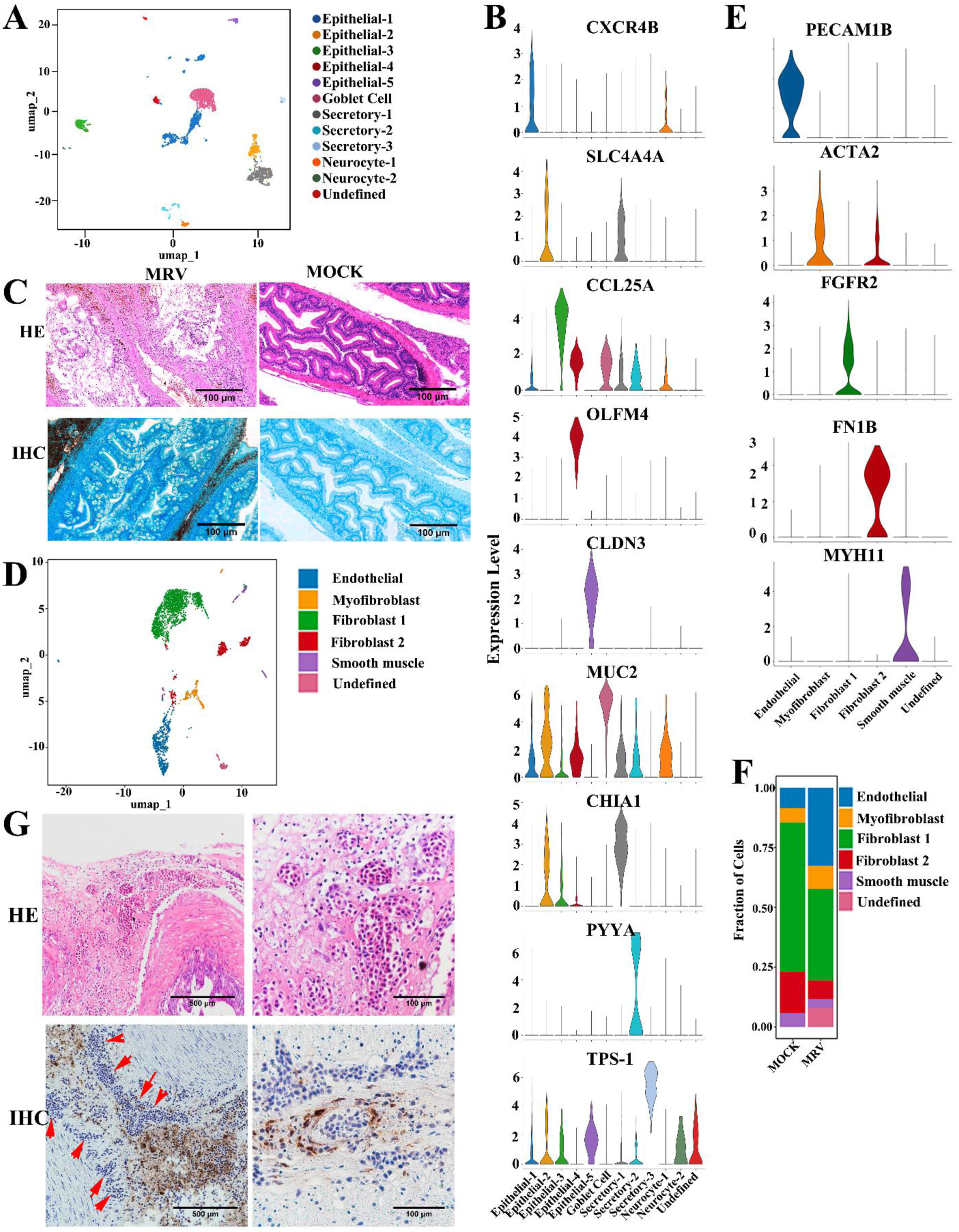
Subtypes of epithelial cells and stromal cells in pyloric caeca. (A) Re-clustering 4,500 epithelial cells of pyloric caecum identified 12 subclusters, shown in UMAP space. (B) Average expression levels and prevalence of major markers used to identify non-secretory epithelial cell and secretory cell of the 12 epithelial cell subtypes. (C) H&E staining and IHC staining of mucosa using anti-MRV mAb. (D) The umap plot shows the 5 stromal cell subtypes identified in pyloric caeca. Different stromal cell subtypes are color-coded. (E) Violin plots showing the average expression level and prevalence of selected major markers used to annotate the immune cell types. (F) Bar plots showing the proportion of each cluster in the stromal cells from control and MRV-infected groups. (G) H&E and IHC of mandarin fish pyloric caeca infection with MRV. The arrows indicate the hypertrophic serosa filled with microvessels. Scale bars, 100 μm.

The global clustering involved 3,410 stromal cells (**Fig. 2B**), which were re-clustered and categorized into six distinct stromal cell types (**Fig. 3D and E**). These cell types include endothelial, myofibroblasts, fibroblast-1, fibroblast-2, smooth muscle and an undefined cell. Among them, there were significant increases in endothelial cells, myofibroblasts, and the undefined stromal cell after MRV infection. In contrast, other cells showed varying degrees of reduction **(Fig. 3F)**. The tSNE plot also showed the expression of selected marker genes which were enriched in subsets (**Fig. S4B**). Through pathological observation and virus tracing in MRV-infected mandarin fish, we once again observed the predominant localization of the virus in serosal layer leading upregulation of numerous genes associated with extracellular matrix (ECM) deposition and remodeling, resulting in severe congestion, edema and hyperplasia (**Fig. 3G and S7A**).

### Compositions and functions of immune cell subpopulations

Upon re-clustering approximately 4,000 immune cells from both infected and healthy mandarin fish, six distinct immune cell subtypes were identified (**Fig. 4A**). Gene expression bubble maps were generated to highlight marker genes for each immune cell subpopulation, and granulocytes, T cells, Macrophages, B cells, NK cells and monocytes, were obtained ((**Fig. 4B, 4C and Fig. S5A**). Among these cell types, T cells and B cells emerge as the predominant immune cell populations in pyloric caeca of healthy mandarin fish (**Fig. 4D**). In response to MRV infection, the numbers of macrophages and granulocytes significantly increased, while T cell numbers significantly decreased. The count of B cells remained largely unchanged (**Fig. 4D**). FISH results also confirmed that numerous ALOXE3^+^ granulocyte and CSF1RB^+^ macrophages/monocyte signals were observed in hypertrophic serosal region (**Fig. 4E and F**). By contrast, few ALOXE3^+^ and CSF1RB^+^ signals were found in the healthy pyloric caecum.

**Fig 4.**
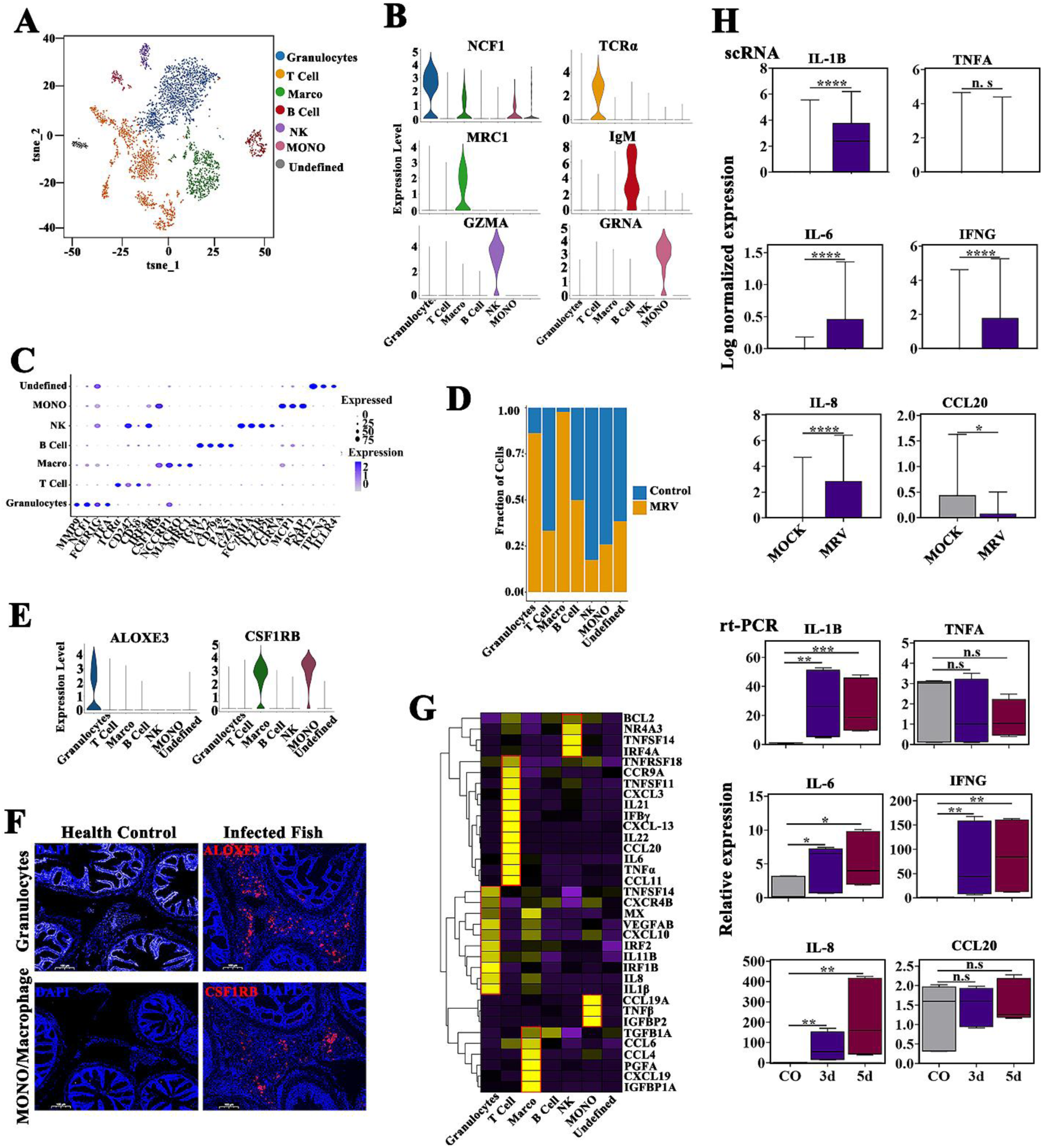
Immunological features of immune cell subsets upon MRV infection. (A) The tSNE plot shows the 7 immune cell subtypes identified in pyloric caeca. Different immune cell subtypes are color coded. (B) and (C) Average expression level and prevalence of selected major markers used to annotate the immune cell types. Violinmap (B) and dotplot (C) of the expression of the marker genes in each immune cell type. (D) Bar plots showing the proportion of each cluster in immune cells from control and MRV-infected groups. (E) Violin plot showing the expression of granulocytes (ALOXE3) and macrophage/monocytes (CSF1RB) marker across the 7 immune cell clusters. (F) FISH data showing the expression of granulocytes subtype marker ALOXE3 (UP) and macrophage/monocytes subtype marker CSF1RB (Down), Scale bar, 100 μm. (G) Heatmap of cytokine expression among 7 immune cell subtypes. (H) Boxplots of cytokine expression based on scRNA-seq and RT-qPCR profiling for healthy control and MRV-infected mandarin fish. Data are expressed as means ± SD. (**p < 0.01, or ***p < 0.001).

To further investigate alterations in gene transcription in immune cells following MRV infection, we compared gene expression patterns between control and MRV-infected cells. Upon viral infection, the expression of most immune-related genes was significantly upregulated. These differentially expressed genes (DEGs) are involved in leukocyte activation and are associated with chemokine signaling pathways (**Fig. S5C**). Subsequently, we examined the inflammatory characteristics of each hyperinflammatory cell subtype to identify potential sources of cytokine production. Distinct pro-inflammatory cytokine gene expressions were identified in each cell subtype (**Fig. 4G**), suggesting that diverse cell types secrete distinct pro-inflammatory factors, contributing to cellular storm phenomena through multiple mechanisms involving cytokine storm induction. We collected scRNA-seq data along with qPCR test results (**Table S1**), both of which confirmed elevated levels of several pro-inflammatory cytokines, such as IL-1β, IL-8, IL-6, and IFNG, in the MRV-infected group. However, not all pro-inflammatory factors exhibited increased expression, including TNFα and CCL20 (**Fig. 4H**).

### Spatial location of infected-pyloric caeca single cells

To further investigate the cellular composition of hyperplastic serosa, the spatial transcriptomics (ST) was employed to map the spatial distribution of scRNA-seq data from MRV-infected pyloric caeca. Analysis of transcriptional signatures of ST spots identified seven spot clusters in the slide, which mapped to discrete anatomical regions (**Fig. 5A and B**). Using the scRNA-seq atlas as a reference, we performed factor analysis to determine the likely single-cell composition of each spot, thus spatially localizing all scRNA-seq clusters. This approach revealed significant differences in cell composition between the gastrointestinal tract, muscle layer, and hyperplastic serosa (**Fig. 5B, C, and S6A**). Furthermore, we observed that the hyperplastic serosa predominantly consisted of fibroblasts, endothelial cells, monocyte/macrophages, T cells, granulocytes, B cells, and erythrocytes (**Fig. 5D, S6B**). Global IF scanning further confirmed that MRV was primarily localized within the hyperplastic serosal region (**Fig. 5E**). These findings underscore the heterogeneity of the hyperplastic serosa and provide insight into why MRV specifically targets this area, rather than the gastrointestinal tract.

**Fig. 5.**
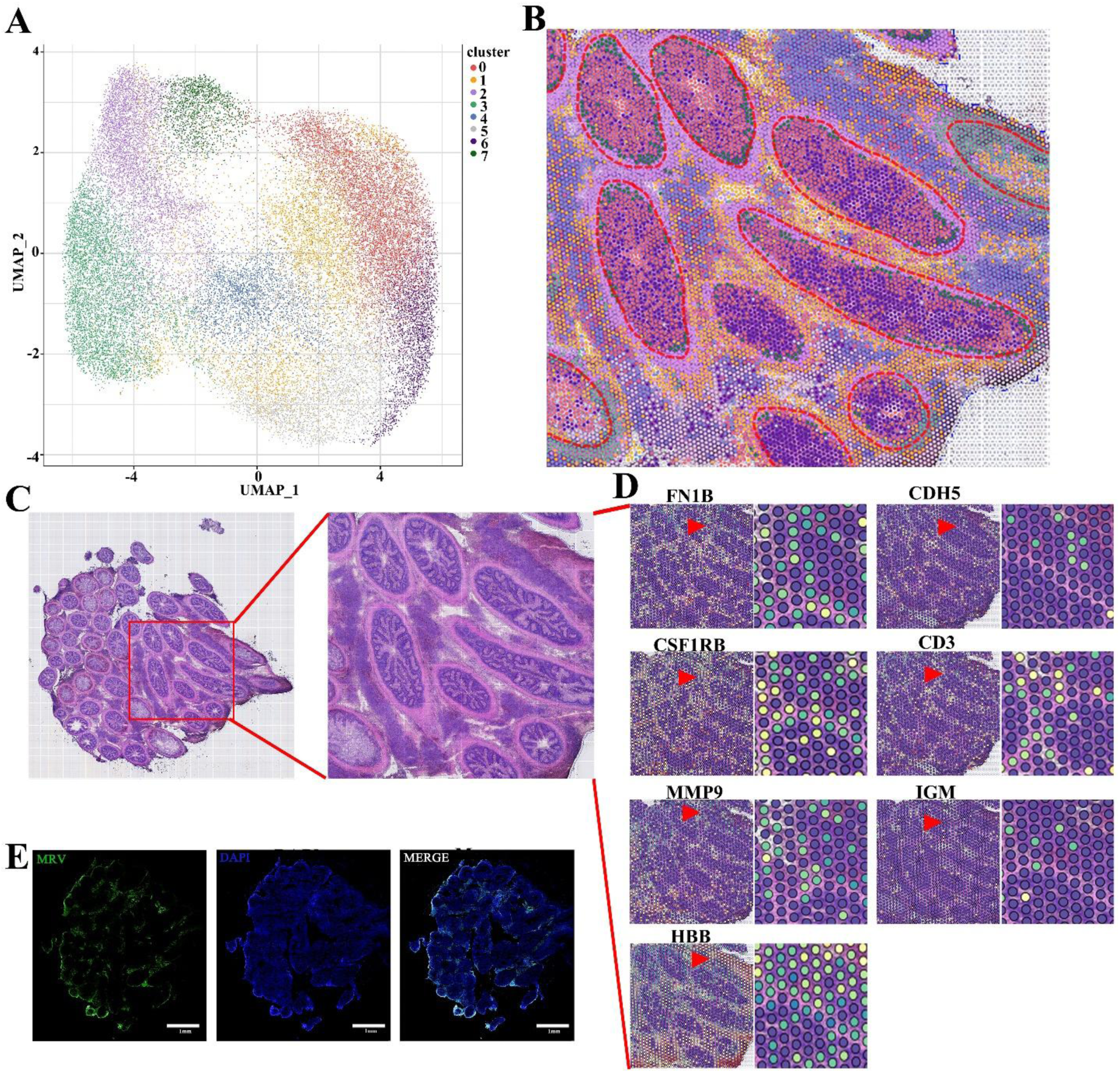
Spatial location of infected pyloric caeca single cells. (A) UMAP plot displaying dimensionality reduction clustering analysis of MRV-infected pyloric caeca by spatial transcriptomics. (B) Spatial transcriptomics data showing the distribution of cell clusters in MRV-infected pyloric caeca in pathological section (partial). The red circle outlines the individual pyloric caeca. (C) Histopathologic microscan of the section of MRV-infected pyloric caeca. The H&E image and the ST are derived from the same tissue section. The red box indicates the enlarged area. (D) Expression of selected cell markers identified by scRNA-seq in hyperplastic serosa. The representative views indicated by the triangular arrows were shown in the enlarged image on the right. (E) IF staining images of MRV (green) in an adjacent section, scale bar, 1 mm.

### MRV mainly attacks B cells, T cells, myofibroblasts, fibroblasts, endothelial and pericytes

The target cells of MRV infection in the pyloric caecum were systematically analyzed using scRNA-seq. Comparison of scRNA-seq data before and after MRV infection revealed a significant increase in the proportion of red blood cells (HBB), epithelial cells (EPCAM), granulocytes (ALOXE3), endothelial cells (CDH5), myofibroblasts (ACTA2), and macrophages (CSF1RB) relative to uninfected cells. Conversely, the proportion of fibroblasts (HSPB1) and T cells (CD3delta) decreased significantly compared to uninfected controls, while the ratio of B cells (IgM) and monocytes remained relatively unchanged (**Fig. 2D and 6A**). Epithelial and muscle cells were excluded from further analysis due to the absence of viral signals in these layers, as observed by IHC and IF (**Fig. 1 and S1**). Dual-color IF analysis revealed the co-localization of MRV signals with HSPB1, IgM, CD3, and ACTA2, indicating that MRV primarily targets fibroblasts, myofibroblasts, T cells, and B cells (**Fig. 6B**). Further studies using FISH and IF identified the co-localization of endothelial cells (CDH5), red blood cells (HBB), macrophages (CSF1RB) with MRV. The results revealed that CDH5^+^ signal was co-localized with MRV signal predominantly within blood vessels (**Fig. 6C**). Vascular structure analysis also confirmed that MRV also infects pericytes (**Fig. 6D**; **Fig. 7B**). No colocalization between ALOXE3^+^, CSF1RB^+^ or HBB^+^ signals and MRV signals were observed (**Fig. 6C**), suggesting that none of granulocytes, macrophages/monocytes or red blood cells were MRV-targeted cells. TEM further supported these findings, revealing the presence of numerous virions in plasma-like cells, lymphocyte-like cells, fibroblasts/myofibroblasts, vascular endothelial-like cells, and pericyte-like cells (**Fig. 6D and Fig. 7B**). In summary, MRV predominantly targets myofibroblasts, fibroblasts, endothelial cells, and pericytes, as well as B and T cells of the infected pyloric caecum.

**Fig 6.**
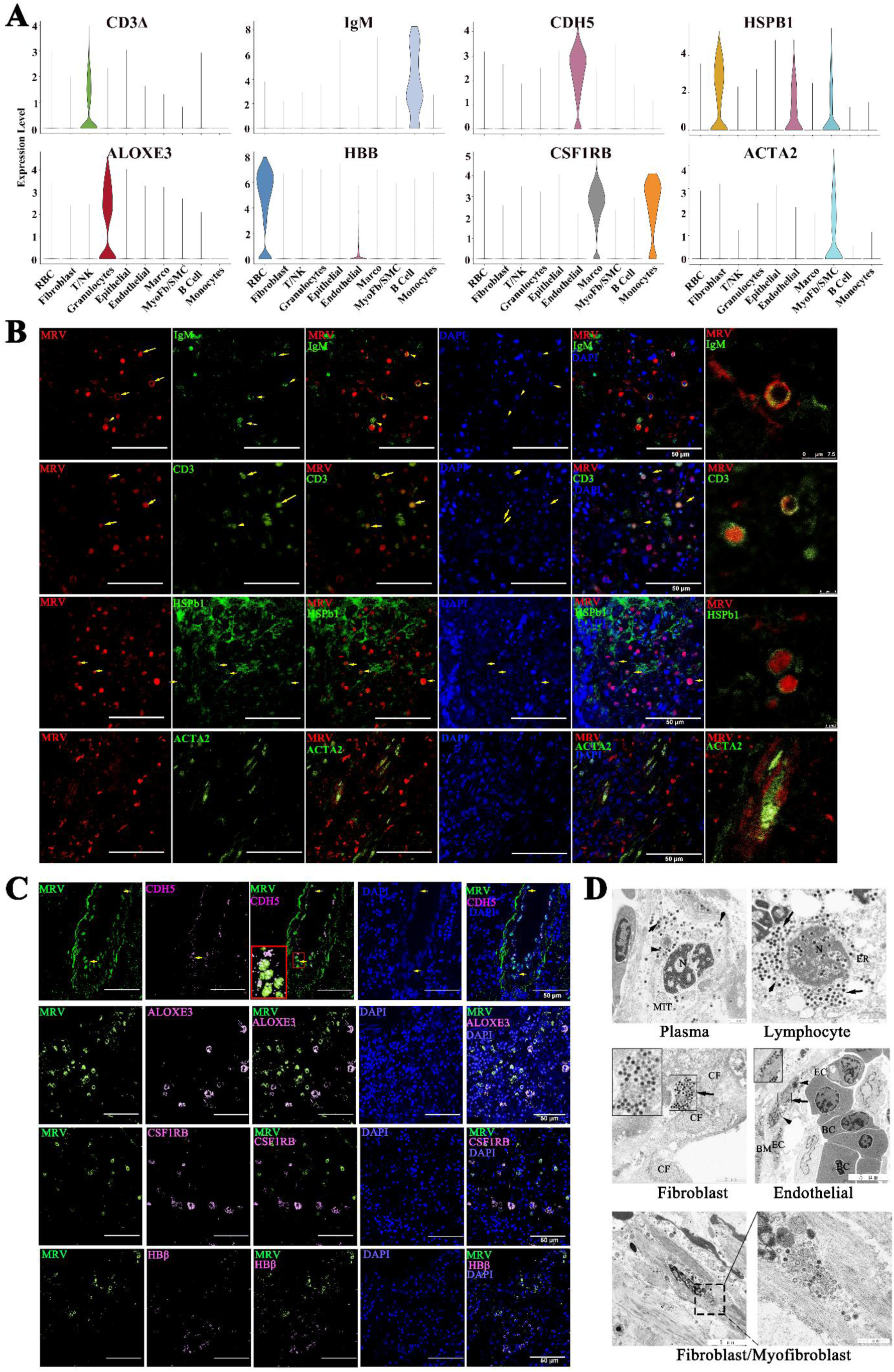
Co-localization of MRV signal and specific markers. Dual-color IF and FISH staining to validate the select marker and MRV signal. (A) Expression levels of selected markers used to identify the selected cell subtypes. (B) Dual-color IF staining to validate the selected markers IgM, CD3, HSPb1, ACTA2 (green) and MRV (red) co-localization. The nucleus was stained with DAPI (blue), Scale bar, 50 μm. (C) Dual-color FISH and IF staining to validate the selected markers CDH5, ALOXE3, CSF1RB, HBβ (pink) and MRV (green) co-localization. Selected markers were identified by the corresponding probe and signal probe (CY5). MRV was identified by anti-MRV monoclonal antibody and secondary goat anti-mouse IgG antibody coupled with AlexaFluor 488 (green). The nucleus was stained by DAPI (blue), Scale bar, 50 μm. The arrows indicate the co-localization cells. (D) Transmission electron micrograph of five kinds of target cell, BC: blood cell, EC: endothelial cell, BM: basement membrane, CF: collagenous fiber, N: nucleus, Mit: mitochondria, ER: endoplasmic reticulum. The arrows indicated the virions. The scale is shown in the figure.

**Fig. 7.**
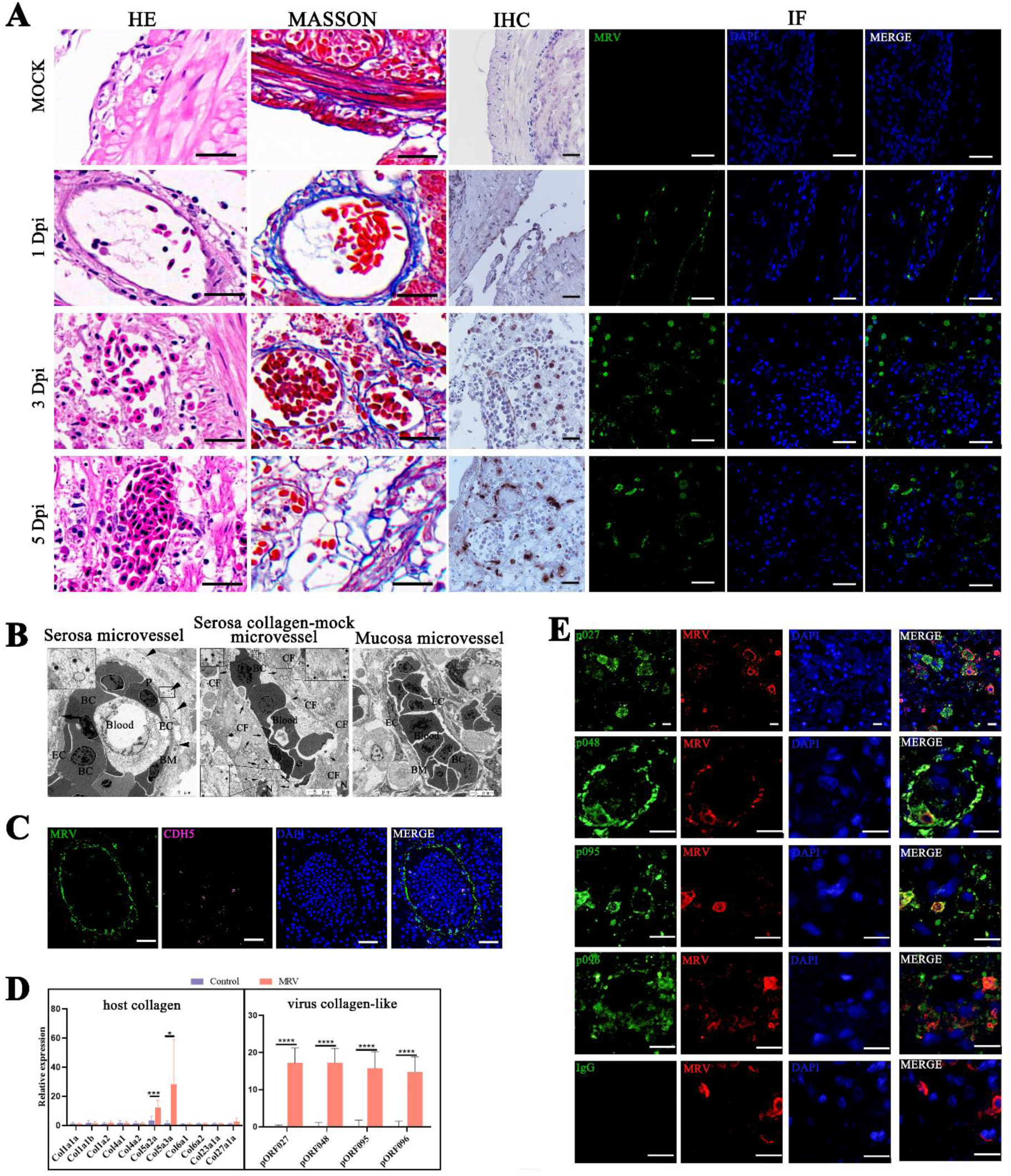
Serosal fibrosis after MRV infection. (A) HE, IHC, IF and MASSON staining observation of pyloric caeca serosa of MRV-infected mandarin fish at 1, 3, 5 dpi, scale bar, 10 μm; black arrows indicated the microvessels; red arrows indicated the MRV virion signals. Scale bar, 20 μm. (B) Transmission electron micrograph of MRV-infected pyloric caeca, (B left) depicts a comprehensive cross-section of microvessels, exhibiting virion-infected endothelial cells and pericytes, scale bar, 2 μm; (B middle) shows a collagen-based vascular lumen devoid of endothelial cells, filled with blood cells and scattered surrounded by virions, scale bar, 5 μm; (B right) showcases intact and distinct mucosal microvessels due to the absence of viral infection in mucosal layer, scale bar, 5 μm. BC: blood cell, EC: endothelial cell, BM: basement membrane, CF: collagenous fiber, N: nucleus. Arrowheads indicated the virions. (C) FISH data showing the localization of virions (green) and CDH5 marker within the microvascular-like lumen. The arrows indicate the shed endothelial cells. Scale bar, 20 μm. (D) The relative mRNA levels of eleven host-encoded collagen (left), and four MRV-encoded collagen-like proteins (right) by RT-qPCR. (E) Co-localization of MRV (red) and collagen-like pORF027, pORF048, pORF095 and pORF096 (green) in infected pyloric caecum by IF. The nucleus was stained with DAPI (blue), scale bar, 10 μm. Data are expressed as means ± SD. (**p < 0.01, or ***p < 0.001).

### Acute MRV infection causes fibrosis of hypertrophic serosa

We identified the target cells of MRV infection through scRNA-seq. Following this, we sought to investigate the underlying mechanisms driving serosal hyperplasia in response to MRV infection. By further analyzing the pathological changes and masson staining of pyloric caeca upon MRV infection, we found that the serosal hyperplasia of pyloric caeca was filled with the vascular lumen, which was mainly composed of collagen fibers **(Fig. 7A)**. With the progress of the infection, the peritoneal structure expanded outward, leading to an increase in the production of collagen fibers and the recruitment of matrix cells and immune cells to combat the viral infection. At the late stage of infection, the interwoven collagen fibers formed a type of extracellular matrix, providing a site for the recruited target cells to adhere **(Fig. 7A)**. Differentially expressed genes (DEGs) associated with extracellular matrix formation, collagen-containing matrices, adherens junctions, and blood vessel morphogenesis were significantly enriched in myofibroblasts, fibroblasts, and endothelial cells in the MRV-infected group (**Fig S7A)**. TEM further revealed two distinct types of microvessels in the hypertrophic serosa: one consisting of normal endothelial cells and pericytes, which were infected by the virus (**Fig. 7B, left**), and the other displaying a vascular lumen composed entirely of collagen fibers. This second type showed a loose lumen structure, devoid of endothelial cells and basal membrane, but accompanied with a significant accumulation of virions (**Fig. 7B middle**). The latter type was more prevalent in the later stages of infection. In contrast, the uninfected epithelial microvessels within the same sample maintained clear structural integrity **(Fig. 7B right)**. FISH confirmed numerous MRV signals around these collagenous microvessels, but only a few endothelial cells were detected, suggesting that these structures are not true microvessels, but collagenous mock vascular structures (**Fig. 7C**).

To investigate the origin of the collagen fibers, we quantified the expression of 11 host collagen genes and observed a significant increase in the expression of Col5a2a and Col5a3a following MRV infection, indicating that the produced collagen fibers were of type V (**Fig. 7D**). MRV whole-genome analysis identified four viral proteins with collagen-like structures, namely pORF027, pORF048, pORF095, and pORF096 (**Fig. S7B**). These viral proteins were highly expressed in the MRV-infected pyloric caeca (**Fig. 7D**). IF analysis showed strong expression of pORF048, pORF095, and pORF096 around collagenous mock microvessels, suggesting their direct involvement in the formation of collagenous structures. Meanwhile, pORF027 was primarily localized in the cytoplasm, indicating potential involvement in other biological processes (**Fig. 7E**). Virion proteome of purified MRV confirmed that these four viral collagen-like proteins are non-structural viral protein. Taken together, these results suggest that MRV actively encodes collagen-like proteins to participate in the formation of the host extracellular matrix, providing a niche for its replication. In the advanced stages of infection, the virus is released from infected cells, and due to the formation of loose fibrous serosal structures, interstitial fluid leaks into the abdominal cavity, leading to ascites formation (**Fig. 1A and Fig. S7C**).

## Discussion

The symptoms of acute MRV infection are systemic. Still, the primary cause of acute mortality in infected mandarin fish is infection-induced hyperemia and edema of pyloric caeca, along with the formation of severe ascites [18, 19]. Investigating the specific site of MRV infection within the pyloric caecum and identifying the targeted cell types are critical for a deeper understanding of the virus’s pathogenesis. To address this issue, we revisited the histopathological changes in the pyloric caecum. Our findings again confirm that the pyloric caeca are the most affected visceral organs during acute MRV infection. Furthermore, the serosal structure of pyloric caecum of mandarin fish was found to be the primary, and possibly the sole site of MRV infection. As the disease progresses, the infected serosa rapidly expands outward, forming a hyperplastic zone that fills the gaps among pyloric caeca. The hyperplastic serosal zone becomes the most densely infected area, leading to hyperemia and edema of the pyloric caeca. We designate this outcome as acute serositis. In LMBV-infected largemouth bass, fibrinous peritonitis, characterization by fibrinous exudates containing numerous leukocytes and abundant cellular debris throughout the peritoneal cavity, is a prominent histopathological feature [13]. Since the peritoneum and serosa are structurally similar, peritonitis or serositis can be considered characteristic histopathological outcomes in both LMBV-infected largemouth bass and MRV-infected mandarin fish. Notably, our recent study proposed that MRV might be a digestive tract virus, linked to ascites syndrome [18]. However, this study clarifies that MRV infects only the serosal structure but not the pyloric caecum cavity itself, indicating that MRV is not a digestive tract virus. This conclusion aligns with the findings in LMBV-infected largemouth bass [13]. In LMBV-infected largemouth bass, exudates are present on the ventral surface of the swim bladder contacting the peritoneal cavity. In contrast, MRV-infected mandarin fish develop severe ascites that fill the abdominal cavity. The mandarin fish’s more developed pyloric caeca [23], compared to the largemouth bass [26], may explain the distinct clinical and histopathological features observed in MRV infection. Overall, irrespective of multi-organ targeted peritonitis or pyloric caecum targeted serositis, the formation mechanism and characteristics of this inflammation response have never been studied. Thus, a comprehensive study based on scRNA-seq was conducted to address this issue.

For scRNA-seq analysis, we identified 10 distinct cell subtypes in the pyloric caeca of mandarin fish, including epithelial cells, five immune cell types, three stromal cell types, and red blood cells. The scRNA-seq profiling revealed a marked enrichment of granulocytes and macrophages in MRV-infected fish. Macrophages, T cells, granulocytes, and NK cells play pivotal roles in the inflammatory response and exhibit a robust inflammatory reaction. Additionally, fibroblasts, myofibroblasts, T cells, B cells, endothelial cells, and pericytes are the primary MRV-targeted cells. Furthermore, MRV infection induces severe fibrotic hyperplasia in serosa, with four collagen-like proteins contributing to the formation of fibrous structures.

Previous studies have extensively characterized the cell types in the gastrointestinal tract of humans and mice [38–40], as well as in the zebrafish intestine [41]. Pyloric caeca, a distinctive and significant structure in fish digestive systems, plays an important role in vertebrate evolution [21,22]. However, knowledge of the cell types in fish pyloric caeca remains limited. In this study, we focused on MRV-infected mandarin fish juveniles (average weight 50 ± 5 g). Single cells from pyloric caeca were collected from two healthy and two MRV-infected fish, yielding 10,777 and 4,807, respectively. Considering the high mitochondrial content in gastrointestinal cells, the threshold for filtering mitochondria was set at 25%. As a result, each control sample contained approximately 2,400 cells, while each infected sample contained over 5,000 cells due to hypertrophic serosa. Additionally, cell type-specific markers were identified to enable precise classification of cell types. Major cellular classes, including epithelial cells, fibroblasts, muscle cells, endothelial cells, and immune cells, could be distinguished based on the expression of category-specific genes associated with their respective functions. In fish, cells from the pyloric caeca can be divided into epithelial cells and other types of cells based on the expression of the pan-epithelial cell marker EPCAM. Epithelial cells were further classified into secretory cells and non-secretory cells based on the secretion of specific proteins. Secretory cell markers are well-defined and functionally distinct. For instance, Secretary-1 expresses chitinase and aminopeptidase, which aid in digesting crustacean food, while EC2 expresses YY peptide. In addition to MUC2-secreting goblet cells, EC3 expresses trypsin1-3 and elastase, key digestive enzymes. Non-secretory epithelial cells lack specific subtype markers, so we categorized them as epithelial-1 to epithelial-5 based on genes that are highly expressed in each subtype. The functions of these subgroups were preliminarily speculated, though further data are needed to refine their classification. Overall, MRV infection induces damage to these epithelial cells; however, these cells are not permissive to MRV infection.

Beyond its digestive function, the gastrointestinal tract plays a crucial role in immunity, with the pyloric caeca of bony fish serving as an key site for immune activity [42]. Immune cells identified in the digestive tract of bony fish include T lymphocytes, B lymphocytes, dendritic cells, macrophages, and granulocytes, all contributing to intestinal immunity [43]. Our scRNA-seq results revealed a rich presence of T lymphocytes (TCRα^+^), B lymphocytes (IgM^+^), granulocytes (NCF1^+^), macrophages (MRC1^+^), NK cells (GZMA^+^), and monocytes (GRNA^+^) in the pyloric caeca of healthy mandarin fish. Among these cell types, T cells, B cells, and NK cells comprised the majority of the immune cell population. Previous studies have shown that peritoneal recruitment and accumulation of NK cells occur three days after ranavirus infection, while lymphocyte recruitment is delayed until 6 dpi [43]. In contrast, our study demonstrated that macrophages and granulocytes exhibited rapid responses to MRV infection, recruiting to the infected site within 3 dpi, while other immune cell types did not show significant enrichment. Non-specific cytotoxic cells (NCC), a fifth type of fish-specific cytotoxic cells and evolutionary precursors of NK cells, were not detected in our study. The marker NCCRP1 specifically expressed in macrophages and granulocytes, whereas granzyme showed specific expression in NK cells. It is possible that NCC cells may have vanished in mandarin fish, suggesting that its immune system may have a higher evolutionary status among bony fish.

In contrast to higher vertebrates, fish are free-living organisms from the early embryonic stages of life, relying on their innate immune systems for survival. Nonspecific immunity is the primary defense mechanism in fish, although their immune memory response is typically less developed than in higher vertebrates [44, 45]. Ranavirus infection is often associated with a prominent host inflammatory response. Similar to viral infections in mammals, ranavirus-induced inflammation is a double-edged sword: while it is essential for viral clearance, it can also exacerbate disease progression and negatively impact host survival [46–48]. Previous studies have observed that ranavirus infection induces the expression of inflammatory genes such as interleukin-1-β (IL-1β), tumor necrosis factor-α (TNFα), and the anti-inflammatory arginase-1 (Arg-1) [43, 49, 50]. Following infection, interferon and the downstream protein Mx are rapidly expressed, though they fail to effectively counter the virus [51, 52]. In this study, we found that the acute MRV infection rapidly caused the increase of many inflammatory factors. Nearly all immune cell types, including granulocytes, macrophages, T cells and NK cells, are involved in the secretion of these inflammatory factors. For example, granulocytes produce IL-1β and IL-8, while T lymphocytes secrete IL-6 and IFNγ, all of which increase significantly within 3 dpi. However, TNFα, secreted by T lymphocytes, does not exhibit significant changes post-MRV infection. These findings suggest that acute MRV infection triggers a strong inflammatory response in the pyloric caeca, aiming to mitigate tissue damage caused by the virus. However, these inflammatory responses are insufficient to clear the virus and may, in fact, worsen the disease and negatively affect host survival. In the case of the pyloric caeca, MRV infection of the serosa leads to severe serositis, characterized by the accumulation of numerous inflammatory cells and the secretion of inflammatory mediators. Additionally, many stromal cells, such as fibroblasts, myofibroblasts, endothelial cells, and pericytes, are recruited to form a hypertrophic serosal zone. Notably, B cells and T cells in the immune cell subpopulation, along with fibroblasts, myofibroblasts, endothelial cells, and pericytes in the stromal cell population, are the primary targets of MRV infection. As a result, MRV infection exacerbates hyperemia and edema within the hypertrophic serosal zone, induces severe ascites, and ultimately leads to high mortality.

The epidemiology, pathogenesis, host antiviral response, and immune escape mechanisms of ranavirus have been extensively studied. However, there have been limited investigations into the specific target cell types of ranavirus infection [53–55]. In chronic infections, very low copy numbers of MRV have been detected in peripheral B lymphocytes, which serve as a reservoir, maintaining a persistently covert infection [53]. Similarly, it was reported that FV3, the type species of the genus *Ranavirus*, can be phagocytic and low copy harbored in macrophages/monocytes [54, 55]. IHC studies have also suggested that the Midwife Toad Virus (a ranavirus) infects liver cells, glomeruli, intestinal mucosa, and endothelial cells of young toads [9, 56]. Despite these findings, global identification of ranavirus target cells remains underexplored. In this study, we revisited and characterized the histopathology of the pyloric caeca in detail. Using single-cell RNA sequencing (scRNA-seq), 18 distinct cell clusters were identified in the pyloric caeca, which were categorized into four groups: epithelial cells, stromal cells, immune cells, and red blood cells. After excluding the epithelial and muscle cells, the remaining cell population includes fibroblasts, myofibroblasts, endothelial cells in stromal group, and B cells, T cells, macrophages, granulocytes, NK cells and monocytes in immune cell group. HIF plus FISH techniques and TEM observations identified fibroblasts, myofibroblasts, endothelial cells, B cells, and T cells as MRV-infected cells. TEM also revealed that MRV infected pericytes, although these cells were not identified in our scRNA-seq results due to the absence of specific pericyte markers. Based on the specific markers (PDGFRB) from other species, pericytes may derive from smooth muscle cells or myofibroblasts. Given that myofibroblasts were identified as MRV target cells (Fig. 6S), it is reasonable to infer that pericytes are also infected by MRV. Interestingly, MRV does not infect macrophages, which is consistent with our recent findings of persistent MRV infection in mandarin fish, but differs from persistent FV3 infection in *Xenopus laevis*, where peritoneal macrophages were confirmed as viral reservoirs [54, 55]. In the mandarin fish model, peripheral B lymphocytes, but not T lymphocytes or macrophages, serve as the viral reservoirs during persistent MRV infection [53]. Persistent MRV infection in fish is characterized by a long-term, low-copy, quiescent MRV state, which can be reactivated by environmental or pharmacological stressors [53]. Notably, T lymphocytes are identified as MRV-targeted cells during acute infection but are non-permissive during persistent infection. We speculate that the robust T-cell-mediated immunity is sufficient to clear the low-copy quiescent MRV during persistent infection. Furthermore, our unpublished data indicate that the attenuated live MRV vaccine, but not the inactivated MRV vaccine, provides effective protection against virulent MRV challenge in the mandarin fish model. This suggests that cellular immunity, rather than humoral immunity, plays a crucial role in combating MRV infection. This may explain why MRV utilizes B cells, rather than T cells, as reservoirs to sustain persistent infections.

Virus infection and dissemination must overcome numerous obstacles, with the extracellular matrix (ECM) of host cells acting as a significant barrier. Some viruses have evolved strategies to bind to specific ECM components to facilitate infection [57]. The ability of viruses to manipulate intercellular communication is crucial for successful infection, as it results in the accumulation of viral particles on the host cell surface, promoting efficient binding to transmembrane or complex host cell receptors [58, 59]. The ECM is a three-dimensional network of macromolecules including collagen, laminin, heparin sulfate, elastin, keratin, chondroitin sulfate, fibronectin, and hyaluronic acid which provide the environment and structure for biochemical support. Studies have shown that integrins, such as α1β1 and α2β1, act as receptors for rotavirus enterotoxins, while integrin α1β1 serves as a cell receptor for Ross River virus [60, 61]. Furthermore, collagen-binding integrin VLA-1 modulates CD8^+^ T cell-mediated immune against heterologous influenza virus infection [62]. Collagen IV (col4a1 and col4a2), induced by human T-cell leukemia virus type 1 (HTLV-1) oncoprotein Tax, forms part of viral biofilms and influences viral transmission [63]. In the present study, we observed that MRV infection of myofibroblasts and fibroblasts in the serosa of mandarin fish pyloric caeca promoted the expression of type V collagen. Additionally, MRV encoded four collagen-like proteins, which, together with host-encoded collagens, formed an ECM network that facilitated the recruitment of target and inflammatory cells. At later stages of infection, the disruption of the ECM and the infiltration of interstitial fluid aided in the rapid release of viral particles following viral replication.

In summary, we established a comprehensive understanding of the pyloric caecum’s response to acute MRV infection in mandarin fish at single-cell resolution (**Fig. 8**). The key finding include, (a) pyloric caecum serves as the most attacked tissue, with the serosa layer bears the primary infection; the acute MRV infection results in hyperplastic serosa and acute fibrinous serositis; (b) The acute fibrinous serositis is characterized by the rapid aggregation of numerous inflammatory cells and the secretion of inflammatory factors; concurrently, a large number of stromal cells such as fibroblasts, myofibroblasts, endothelial cells and pericytes were recruited to form hypertrophic serosal zone; (c) T cells, B cell, fibroblasts, myofibroblasts, endothelial cells and pericytes in the hypertrophic serosal zone are targeted by MRV, driving ECM formation, collagen encoding, and blood vessel morphogenesis; the infected pyloric caeca ultimately shows congestion, edema, and adhesions; (d) the loose ECM and the interstitial fluid infiltration facilitate the rapid release of the virus, further exacerbate the progression of serositis, leading to severe ascites and mortality. One limitation of this study is the relatively small number of biological replicates used for scRNA-seq, which constrained the depth of analysis and exploration of the single-cell data. This limitation stems from the lysis of cells during the terminal stages of viral infection (**Fig. S7C**), which results in the loss of some virus-infected cells during suspension preparation and may influence statistical analyses. Nevertheless, these limitations do not compromise the robustness of our conclusions regarding the identification of viral target cells or the mechanisms by which infection drives serositis.

**Fig. 8.**
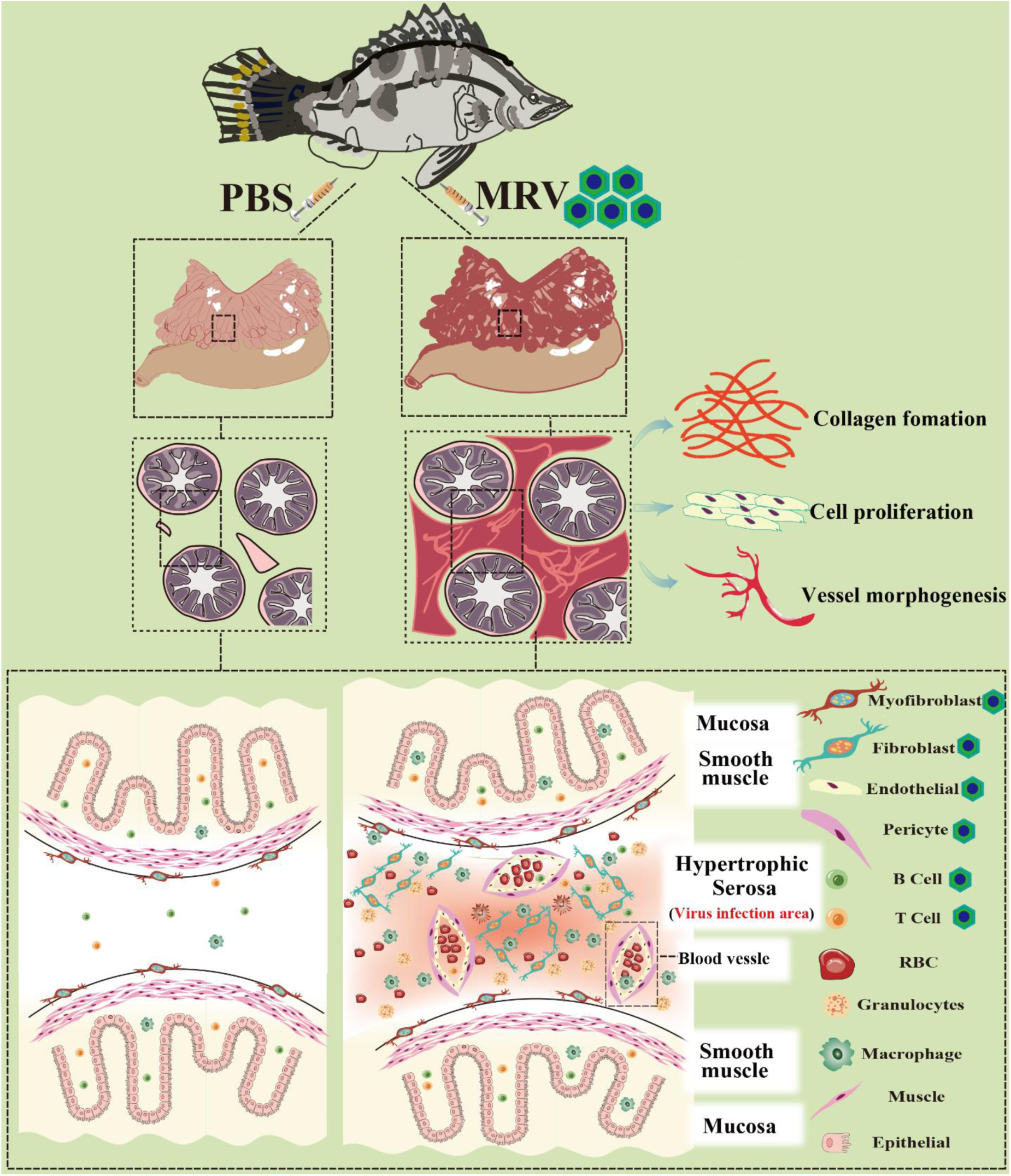
Diagram illustrating pyloric caeca lesion in mandarinfish upon acute MRV infection. The acute MRV infection results in severe fibrinous serositis characterization by severe congestion and edema in pyloric caeca, which is attributed to MRV primarily infecting mandarin fish serosa and resulting in pronounced serosal hyperplasia. Furthermore, MRV predominantly infected stromal components such as myofibroblasts, fibroblasts, endothelial cells, pericytes, and T and B lymphocyte within immune cells. This led to rapid ECM formation and recruitment of stromal and immune cells in hypertrophic serosal zone, creating a favorable environment for viral replication. Simultaneously, due to the loose ECM structure, significant cell infiltration and interstitial fluid leakage resulted in severe ascites syndrome.

## Materials and methods

### Virus, cells and antibodies

MRV-ZQ17 strain was isolated and characterized from a natural mass mortality of mandarin fish juvenile and propagated in mandarinfish fry (MFF-1) cell line [18]. The MFF-1 cell line was grown in complete Dulbecco’s modified Engle’s medium (DMEM) (Gibco, USA) containing 10% fetal bovine serum (FBS) (Gibco, USA), at 26 °C containing 5% CO_2_ [64]. Monoclonal antibodies (mAbs) specific to MRV 1C4 and mandarin fish IgM were constructed, characterized and stored in our laboratory [19]. The rabbit polyclonal antibodies of anti-MRV-027R, anti-MRV-048L, anti-MRV-095R, anti-MRV-096R, and anti-mandarin fish T-cell surface glycoprotein CD3delta were customized by GL Biochem (China). Additional rabbit polyclonal antibodies, including anti-actin alpha 2 (ACTA2) and anti-heat shock protein family b (small) member 1 (HSPB1), were obtained from ABMART company (China).

### Animals and artificial infection

Juvenile mandarin fish (average weight: ∼45 g) were sourced from a fish farm in Foshan, Guangdong Province, China. All animal experiments were conducted in accordance with the animal testing procedures of Guangdong Province, China, and were approved by the Ethics Committee of Sun Yat-sen University (Approval number: SYSU-LS-IACUC-2024-0016). The fish were confirmed to be free of MRV and ISKNV via qPCR as previously described [19], and divided into two groups, 30 fish per group. One group was used for MRV infection, while the other served as the mock-infected control. The water temperature was kept at 26 ± 0.5 °C. For the infection group, an MRV-ZQ17 suspension cultured in MFF-1 cells was adjusted to a concentration of 10^7^ TCID_50_/0.1 mL and administered by intraperitoneally (i. p) injection with 0.1 mL per fish. The control group received an equivalent volume of sterile PBS. At 1-, 3- and 5-days post infection (dpi), both infected and control fish were sampled for quantitative gene expression analysis, tissue tropism studies and sc-RNA seq.

### Sample collection and dissection of scRNA-seq

Four pyloric caecum samples were collected, comprising two from MRV-infected fish and two from healthy controls. Single-cell suspensions were prepared from tissue fragments using a combination of enzymatic and mechanical dissociation. Cell capture and cDNA synthesis were carried out on a 10×Genomics platform. Transcriptome sequencing was performed on a NovaSeq 6000 platform after meeting all quality controls.

### Processing scRNA-seq data

Raw data generated with the 10× Genomics platform was aligned to the reference genome using Cell Ranger software (v7.0.1) to obtain the UMI matrix, which was further imported into R (v4.2.2) and processed with the Seurat package (v4.3.2). Cells with a detected gene number <200 or >5000 or a high mitochondrial transcript ratio (>25%) were excluded. After normalization and scaling, the batch effect between patients was then removed using harmony. The batch-corrected matrix was used for further analysis and visualization. The top 2000 highly variable genes were extracted to perform principal component analysis (PCA), and the top 30 principal components (PCs) were used for cluster analysis. Cell type was annotated by the SingleR package (v1.2.4) and then checked manually.

### Identification of marker genes

To identify differentially expressed marker genes for each cell type, the Find All Markers function in Seurat was used under default parameters. Marker genes were selected as those with adjusted p values less than 0.05, average logFC larger than 0.25, and percentage of cells with expression higher than 0.25. Gene Ontology (GO), Kyoto Encyclopedia of Genes and Genomes (KEGG) and GSEA were performed with the R package, cluster Profiler v3.18.0, using all detected genes from the entire scRNA-seq library as background. Terms were enriched with the nominal p value < 0.05 and false discovery rate (q value) < 0.05.

### Copy-number analysis

InferCNV (version 1.2.3; ref. 21) was used to infer large-scale copy number variations in immune cells using count data as input. Filtering, normalization, and centering by normal gene expression were performed using default parameters and data was scaled. A cutoff of 0.1 was used for the minimum average read counts per gene among reference cells. An additional denoising filter was used with a threshold of 0.2. Copy-number variation was predicted using the default six-state hidden Markov model.

### Sample preparation, library construction, sequencing and data analysis of ST

Fresh MRV-infected pyloric caecum was frozen using isopentane and liquid nitrogen bath; The frozen tissue samples were embedded with OCT and stored at -80 °C. The frozen slides were carried out in the cryo- ultramicrotomes, and slides were collected for RNA extraction for quality control. RNA integrity number (RIN) values of ≥7 were required to confirm that RNA remained intact during tissue freezing. Tissue sections were mounted onto gene expression slides, followed by methanol fixation, hematoxylin and eosin (H&E) staining, and bright-field imaging. Tissue permeabilization was performed based on the permeabilization time determined during tissue optimization. Released mRNA from the tissue sections was captured by primers on the slides and reverse-transcribed into cDNA; the cDNA was collected from the slides and subjected to second-strand synthesis, denaturation, and PCR amplification. The amplified cDNA was fragmented enzymatically, end-repaired, A-tailed, and purified using beads. Fragments were screened, adapters ligated, and purified again before indexing through PCR to construct a standard next-generation sequencing (NGS) library. Qualified libraries were sequenced using an NGS platform in paired-end 150 (PE150) mode. Sequencing data were processed using BSTMatrix software (Biomarker tech., China) for BMKMANU S1000 spatial transcriptome sequencing. The likely single-cell composition of each spot was determined by carrying out factor analysis using scRNA-seq atlas as a reference, thus spatially localizing all scRNA-seq clusters.

### Viral load measurement by absolute qPCR

The MRV genome copy number was determined by absolute qPCR. DNA templates were extracted from different mandarinfish tissues at 5 dpi using FastPure Cell/Tissue DNA Isolation Mini Kit (Vazyme, China) according to the manufacturer’s instructions. An absolute standard curve of the MRV *mcp*-specific qPCR system was established through diluted pMD-19T-MCP recombinant plasmid DNA. The reactions were performed using the Light Cycler® 480- II Multiwell Plate 384 real-time detection system (Roche Diagnosis, the USA) under the following conditions: 1 cycle at 95°C for 60 s, and 40 cycles at 95°C for 5 s, 60°C for 30 s and 70°C for 5 s with a total reaction volume of 10 μL, containing 1 μL of DNA, 5 μL of Real Time PCR Easy^TM^-Taqman (Foregene , China), 0.5 μM primers, 0.2 μM probe (Tsingke, China) and 3 μL of RNase free water. All real-time qPCRs were performed in triplicate.

### Quantitative gene expression analyses

Total RNA from various tissues of infected mandarin fish at 0-,1-, 3-, and 5-dpi were extracted using Eastep Super Total RNA Extraction Kit (Promega, China) according to the manufacturer’ s instruction. RNA was reverse-transcribed into cDNA using Evo M-MLV RT Premix for qPCR (Accurate Biology, China). The gene expression levels were quantified by qRT-PCR methods in Roche LightCycler 480 system, with mandarin fish β-actin as reference gene. Gene-specific primers were designed and validated by gradient PCR and with the qPCR melting curves. All primers used in this study are listed in Table S2. The qRT-PCR was performed as described previously [18]. Briefly, PCRs were performed as the following procedure: 95 °C for 5 min, 1 cycle; 95 °C for 5 s, 60 °C for 30 s and 70 °C for 5 s, 40 cycles; 95 °C for 5 s, 60 °C for 1 min and 95 °C, 1 cycle; 50 °C for 30s, 1 cycle with a total reaction volume of 10 μL, containing 1 μL of cDNA, 5 μL of 2 × SYBR Green Pro Taq HS Premix (Accurate Biology, China), 0.5 μM primers (Tsingke, China) and 3.6 μL of RNase free water.

### Histopathology, immunofluorescence assay (IFA) and immunohistochemistry (IHC)

Tissues from three moribund fish were dissected and fixed with alcohol-formal-acetic (AFA) for hematoxylin-eosin (H&E) staining, or in 4% paraformaldehyde for IFA and IHC analysis. The fixed tissues were embedded and excised into sections, dewaxed in xylene, and rehydrated in a series of ethanol solutions, according to protocols described in our previous reports [18]. For IHC analysis, the sections were performed using mouse anti-MRV mAb of 1C4 (1:1000) as the primary antibody and horseradish peroxidase (HRP)-labelled Goat anti-mouse IgG as the second antibody, then developed with 3,3 N-diaminobenzidine tertrahydrochloride (DAB) solution. Sections were visualized under a Nikon fluorescence microscope (Eclipse Ni-E, Japan). For simple or double staining IFA (**Fig. 1, 5, 6B, 7A and 7E**), the sections were stained with primary anti-MRV monoclonal antibody and corresponding polyclonal antibody at 1:1,000 dilutions and secondary Alexa Fluor 555-labeled monkey anti-mouse IgG antibody and Alexa Fluor 488-labeled goat anti-rabbit IgG antibody at 1:1,000 dilution, then stained by 4′,6- diamidino-2-phenylindole (DAPI) (Abcam, China). For dual- anti-mouse double staining IFA **(Fig. 6B-IgM**), the sections were stained with the Three-color Multiplex Immunohistochemical Kit (Shanghai YEPCOME Biotechnology Co., Ltd, China) according to the manufacturer’ s instruction. Finally, sections were visualized under a confocal laser scanning immunofluorescence microscopy (Leica SP8, German).

### Fluorescent in situ hybridization (FISH) and IFA

FISH sweAMI riboprobes and corresponding branch probe targeting the open reading frame (ORF) sequences of *CDH5*, *ALOXE3*, *CSF1RB*, *HBβ* and *EPCAM* genes were customized from Servicebio (Servicebiotech, China). The riboprobe sequences are provided in Supplementary S1 (Table S1). Pyloric caeca were rinsed with PBS, and immediately put into in-situ hybridization fixative (Servicebiotech, China) for over 12 h, and stored at 4 °C. After fixation, the target tissue blocks were sectioned to approximately 3 mm thickness under a fume hood, dehydrated through a graded ethanol series, and cleared with xylene. The tissue blocks were embedded in paraffin and sectioned into 4 μm slices, which were baked at 62 °C for 2 hours. After dewaxing, dehydration and antigen retrieval, pre-hybridization at 37 °C for 1h, the sections were hybridized overnight in a constant temperature chamber. After hybridization, sections were sequentially washed twice in 2×saline-sodium citrate (SSC) (1×SSC = 0.15 M NaCl, 15 mM Na citrate) at room temperature for 15 min, then in 1×SSC and 0.1×SSC at 55 °C for 1 h. Branch probes were applied for hybridization at 40 °C for 45 minutes. After washing, CY5-labeled signal probes were added and incubated at 42 °C for 3 hours. Following FISH, tissue sections were stained with primary anti-MRV monoclonal antibody, followed by secondary Alexa Fluor 488-labeled monkey anti-mouse IgG antibody at 1:1,000 dilution, then stained by 4′,6- diamidino-2-phenylindole (DAPI) (Abcam, China). Sections were visualized under a Nikon fluorescence microscope (Eclipse Ni-E, Japan).

### Transmission electron microscopy (TEM)

TEM analysis was performed as our previous description [17]. MRV-infected pyloric caeca from moribund mandarin fish at 5 dpi were collected for TEM assays. Briefly, infected tissues were fixed with 2.5% glutaraldehyde in 0.1 M PBS, followed by secondary fixation in 2.0% osmium tetroxide in 0.1 M PBS. After dehydrating, penetrating, embedding, and polymerization, ultrathin sections (60 nm) were prepared. Sections were stained with uranyl acetate-lead citrate and examined under a Philips CM10 electron microscopy.

### Statistical analysis

Analysis of variance (ANOVA) was performed for statistical analysis of expression and viral load data. Statistical analysis of survival data was performed using a Log-Rank Test (GraphPad Prism 8, San Diego, CA, USA). * means p < 0.05; ** means p < 0.01, *** means p < 0.001, **** means p < 0.0001, ns means no significance. Error bars on all graphs represent the standard error of the mean (SEM).

### Supporting information

**Fig S1. Temporal changes of infected pyloric caeca during acute MRV infection.** (Left), Histopathological changes of infected pyloric caeca visualized by H&E staining at 1, 3 and 5 dpi, showing progressive serositis classified as slight, mild, and severe. The interface between healthy pyloric caecum is clear, with minimal mesenchymal infiltration. (Middle), IHC analysis illustrating MRV distribution in the pyloric caeca over the infection timeline. (Right), IF imaging tracing the progression of MRV infection in the pyloric caeca. Scale bars are shown in the figure.

**Fig. S2 Data quality control of scRNA-seq.** (A) Distribution of gene number detected in single cell across 4 samples (Y axis). (B) Distribution of the percentage of mitochondrial gene expression in single cell across 4 samples (Y axis). (C) Relationship between the number of nUMI and nGene. (D) Relationship between the number of nUMI and pMito.

**Fig. S3 UMAP and violin plots illustrating the expression of representative marker genes.** UMAP and violin plots show that the expression of representative marker genes is restricted to specific clusters among all cells.

**Fig. S4. UMAP plots illustrating marker gene expression in epithelial and stromal cell clusters.** (A) UMAP plots showing that the expression of representative marker genes is restricted to specific clusters within epithelial subtype cells. (B) UMAP plots showing that the expression of representative marker genes is restricted to specific clusters within stromal subtype cells.

**Fig. S5 the expression of representative marker genes and the DEGs of immune cells.** (A) Umap plots showing that representative marker genes are restricted to specific immune cell clusters. (B) tSNE plots aligning clusters of immune subtype cells between control and MRV-infected mandarin fish. (C) Volcano plot illustrating DEGs in immune cells between control and MRV-infected groups. Genes specifically upregulated or downregulated are highlighted in red and blue, respectively. Each dot represents an individual gene, with significant genes identified by adjusted p < 0.05 (adjusted by false discovery rate in MAST). Non-significant genes are shown in gray.

**Fig. S6. Spatial transcriptomics analysis of MRV-infected pyloric caeca.** (A) Spatial transcriptomics data showing the distribution of cell clusters in pathological sections of MRV-infected pyloric caeca. (A) Expression of selected cell markers identified by scRNA-seq in MRV-infected pyloric caeca.

**Fig. S7** (A) Comparison of GO terms for MRV target cells between control and MRV-infected mandarin fish, labeled with names and IDs and sorted by −log10 (P) value. (B) The top 20 enriched GO terms are shown. (B)Prediction of four MRV-encoded collagen-like protein domains using SMART. (C) TEM images showing the lytic cells at the end stage of infection. Arrows indicated the MRV virions.

**Table S1** The underlying numerical data for Fig 4H.

**Table S2** Primers used in the RT-qPCR and the sequence used for synthesis SweAMI FISH riboprobes.

## ACKNOWLEDGMENTS

This work was funded by Guangdong Basic and Applied Basic Research Foundation under No. 2023A1515010009; The key areas R&D Program of Guangdong Province under No. 2021B0202040002; Innovation Group Project of Southern Marine Science and Engineering Guangdong Laboratory (Zhuhai) under No. 311021006.

